# Leaders are made: Learning acquisition of consistent leader-follower relationships depends on implicit haptic interactions

**DOI:** 10.1101/2021.12.09.471486

**Authors:** Asuka Takai, Qiushi Fu, Yuzuru Doibata, Giuseppe Lisi, Toshiki Tsuchiya, Keivan Mojtahedi, Toshinori Yoshioka, Mitsuo Kawato, Jun Morimoto, Marco Santello

## Abstract

Are leaders made or born? Leader-follower roles have been well characterized in social science, but they remain somewhat obscure in sensory-motor coordination. Furthermore, it is unknown how and why leader-follower relationships are acquired, including innate versus acquired controversies. We developed a novel asymmetrical coordination task in which two participants (dyad) need to collaborate in transporting a simulated beam while maintaining its horizontal attitude. This experimental paradigm was implemented by twin robotic manipulanda, simulated beam dynamics, haptic interactions, and a projection screen. Clear leader-follower relationships were learned despite participants not being informed that they were interacting with each other, but only when strong haptic feedback was introduced. For the first time, we demonstrate the emergence of consistent leader-follower relationships in sensory-motor coordination, and further show that haptic interaction is essential for dyadic co-adaptation. These results provide insights into neural mechanisms responsible for the formation of leader-follower relationships in our society.

---

A distinctively unique feature of human culture is the creation of social institutions, defined as sets of behavioral practices that are regulated by different types of mutually recognized norms and rules^1^. Throughout the evolution of human societies, rules have helped define the roles of leaders in a group, i.e., chiefs and presidents. These leaders, in turn, establish rules or norms for the group – i.e., followers – to adhere to so as to ensure that common goals can be achieved and benefit the entire community^2^. Thus, *cooperation* emerges as organized and agreed-upon ways of interacting among members of the group. The cognitive processes underlying cooperation are known as *collective intentionality*^3^, through which the ability of creating joint intentions and commitments in cooperative actions emerges^1^. Therefore, for cooperation to succeed, necessary conditions include a mutual understanding of a common goal, interaction rules that all cooperating agents are expected to follow, and the prioritization of attaining a common – rather than individual – goal. It is generally acknowledged that dyadic interactions are a fundamental unit of large-scale interactions among many agents^2,4^. Therefore, dyadic interactions have been extensively used as a model to understand the mechanisms responsible for large-scale human-human cooperations.

One type of dyadic cooperation is mediated by motor activity by which two agents interact to perform a task to attain a common goal. It has been often observed that leader-follower relations can emerge even when in absence of a priori role assignment^5–7^, which may manifest in task-specific ways. In some tasks, the leader temporally leads the actions, while the follower lags behind^8–10^. In other tasks, the leader may exhibit more corrective behavior and variability^11–14^. Further, in tasks where agents are connected by a physical medium, dyads exhibit asymmetrical sharing of forces^15–17^ and subject-specific joint coordination patterns^18^. However, why and how a particular coordination asymmetry emerges remains unknown^19^. For example, each dyad performing a rhythmic joint tracking task can consistently adopt one of many strategies, even though there was no convergence to a global strategy for agents to share their contributions^20^. Furthermore, the appearance of ‘roles’ may not necessarily mean that they were chosen purposefully through dyadic decision making. In some cases, asymmetrical contributions may be just a by-product of mismatch in individuals’ sensorimotor control capabilities^16,21^. It is conceivable, however, that asymmetrical contributions might also emerge to fullfil a functional goal, e.g., to improve performance or minimize effort. In both of these instances, however, the mechanisms underlying dyadic co-adaptation have not been addressed in the literature.

To further our understanding of role emergence and its functional role in motor interactions, we designed a novel collaborative task in which two agents moved a dynamically simulated, visually-rendered beam through concurrent motion of their hand via a pair of robotic manipulanda (**Fig. 1a**). Each subject was instructed to move the beam to a target (10 cm away from the start position) while not exceeding the maximum tilt (±1.15 degrees) within no shorter than 1 but no longer than 2 seconds (**Fig. 1b**). The agents were located separately and they could not exchange verbal cues nor see each other during task performance. Note that although subjects were told that they were going to execute the tasks on their own, neither subject can complete the task with a single hand because the beam tilt would become too large. Furthermore, subjects were unaware of underlying asymmetry of the task, which was implemented by each agent acting on the beam at different distances from a translational pivot point (**Fig. 1c**). Importantly, the pivot and hands movement were constrained only in the y-direction by a virtual frictionless rail. Unlike previous work using symmetrical dyadic tasks, our asymmetrical task inherently provides gradients for optimization such that some coordination strategy has certain advantage over others. We hypothesized that this gradient for optimization of performance variable(s), e.g., beam angle, would drive dyads to co-adapt towards an optimal strategy that is consistent across dyads. Based on previous work^22–24^, we further hypothesized that dyads would adapt slower in absence of haptic feedback. This hypothesis is based on the expectation that reliable information exchange between the participants is a pre-requisite for dyadic co-adaptation.

**Figure 1.**
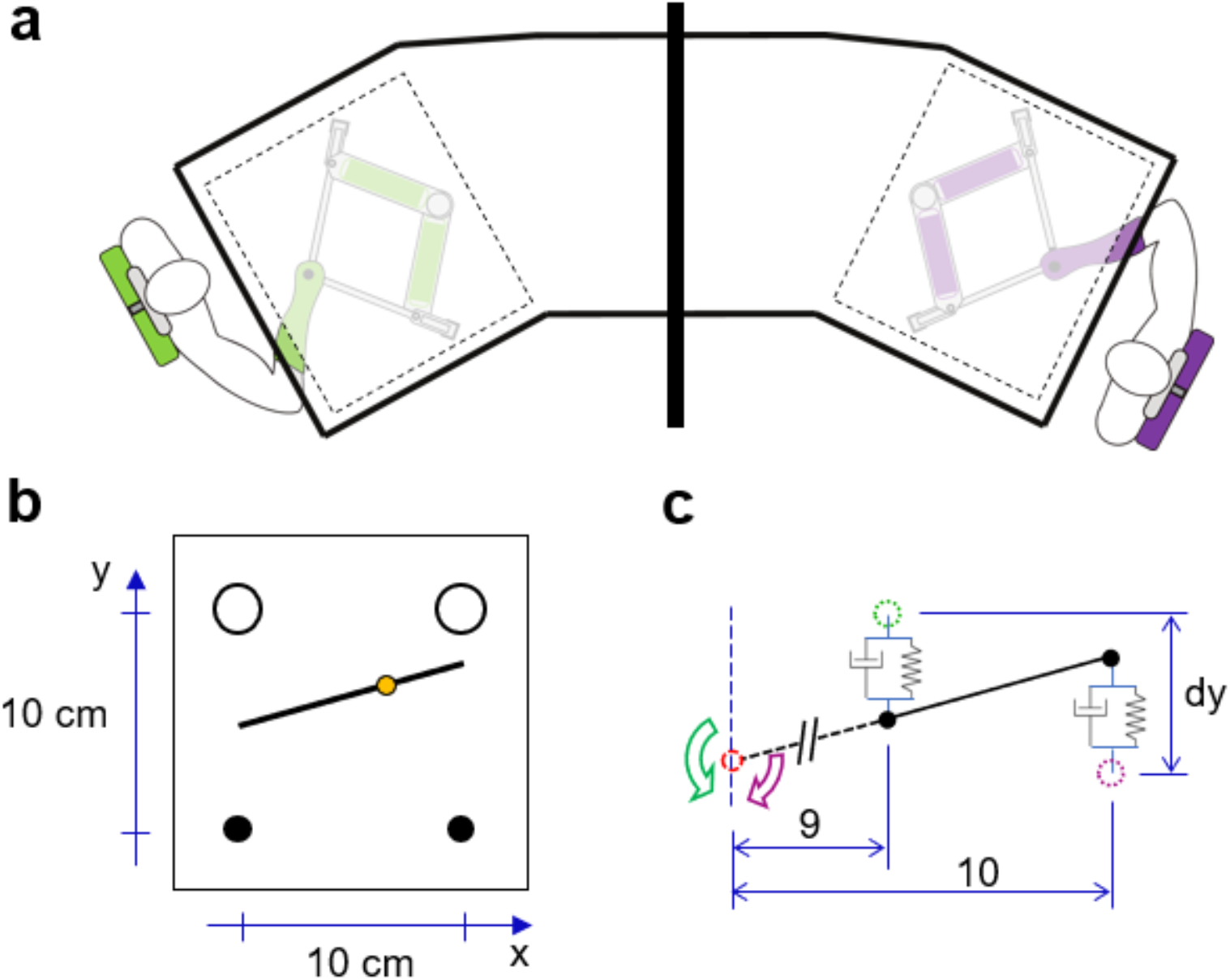
Experimental setup. **a** Each subject grasped and moved a robotic manipulandum located under a table. The two manipulanda were used to simulate the physical interaction between two human subjects. The virtual beam linking the two manipulanda was displayed on the table. The actual hand position was blocked from view. Visual feedback of the partner was blocked by black boards placed between subjects. Subjects wore headphones to eliminate sounds generated by the partner. Dyads performed the task without being informed that they were interacting through the virtual beam. **b** Visual feedback of the physical interaction task. Subjects were asked to move a 10 cm long and 1 kg weight virtual beam from the start position to the end position (bottom and top circles, respectively; 10 cm distnace) while maintaining the beam horizontal. The tilt indicator (yellow circle) behaves like an air bubble in a level scale and slides towards the higher of the two beam’s ends. **c** Task dynamics. The beam behaves as if a pivot is located at one side of the beam far away from the hands (pivot to hand distance ratio between right and left agent was 9:10). Each hand generates a force to move the beam through a virtual spring-damper system between the hand position (dashed circle) and corresponding beam attachment position (black dot).

## RESULTS

The beam’s movement in our task was dependent on the coordination of the hand movement controlling the left and right side of the beam. The parameters of the task was selected to afford a large space of coordination strategies. One metric that quantifys the movement coordination is to compute the mean hand position difference, which indicates the spatial relation between two hands. Computer simulation using minimum jerk hand kinematics suggested that the mean hand position difference (Left – Right; Fig. 1C, dy) could be within a range of -0.02 m to 0.06 m while complying with the task’s spatial and temporal requirements (see simulation results in **Supplementary Fig. 1**). This large range indicates that the task is relatively easy to perform successfully, and either hand could be spatially advanced with respect to the other hand. As expected, participants were able to accomplish 40 successful trials across all stiffness conditions with only several failed trials (**Supplementary Fig. 2a**). Movement speed increased across successful trials in all conditions. All dyads started at a similar movement speed in the first five successful trials (completion time: 1.41 ±0.19 s) and ended with similar speed in the last five successful trials (completion time: 1.23 ±0.20 s). Two-way mixed ANOVA revealed only a significant main effect of Trial (F(1,21) = 21.54, *p* < 0.001, **Supplementary Fig. 2b**), but not Stiffness.

Importantly, we observed that dyads in S400 condition, in which high stiffness haptic feedback was provided, clearly exhibited systematic and consistent changes of motor behavior across successful trials (**Fig. 2a**). In contrast, dyads in S0 condition, who had only visual feedback, only showed random and highly variable trial-to-trial changes (**Fig. 2b**). This difference is evident in rainbow-like color distributions in **Fig. 2a** and **c** versus the intermingled color distributions in **Fig. 2b** and **d**. Specifically, consistent spatial and temporal relation emerges in S400 with the movement of the agent controlling the left side leading the right side. Moreover, the magnitude of the left spatiotemporal leadership gradually increase as more trials were completed, which is evident from clear color gradients across position and force trajectories in (**Fig. 2a** and **c**). A clear emergence of asymmetrical role can also be observed in the force measured at the robotic handle in S400 condition (**Fig. 2c**), but not in S0 (**Fig. 2d**). The left agents gradually increased the forces in the forward direction as their spatiotemporal leadership grew, while the right agent gradually increased the force in the backward direction. Furthermore, we found that the left agent exhibited a larger variation in the force they produced than the right agent, suggesting a large extent of corrective behavior.

**Figure 2.**
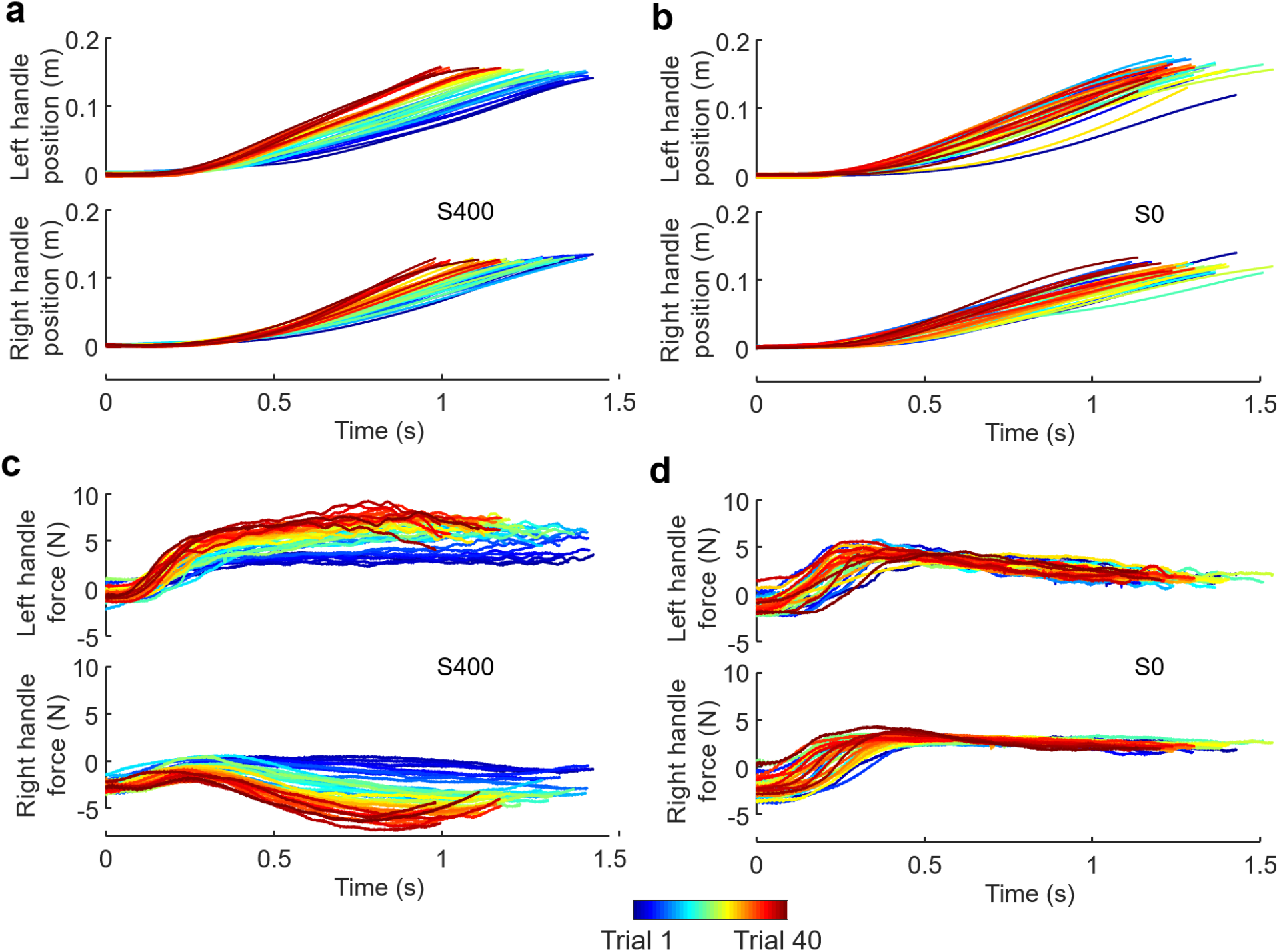
Changes in motor behavior of paired individuals across successful trials. Early trials are plotted as blue traces, and subsequent trials are denoted by ‘warmer’ colors. **a** Left and right handle movements performed by a representative dyad in the S400 condition. **b** Hand movement performed by a different representative dyad in the S0 condition. **c** Measured left and right handle force from the same dyad shown in **a. d** Measured handle force from the same dyad shown in **b**. Rainbow-like color distributions in **a** and **c** clearly indicate consistent changes of behavior across trials in the S400 condition, whereas intermingled color distributions in **b** and **d** indicate no such consistent changes for the S0 condition.

Statistical comparisons supported the above observations. We first quantify the spatial relation between the paired agents by computing the hand position difference, such that a positive value would suggest spatial lead of the left agent. As indicated by the trajectories of individual agents (**Fig. 2**), we confirmed a consistently increasing spatial lead of the left agent in S400 condition (**Fig. 3a**) but not S0 condition (**Fig. 3b**). For the medium stiffness condition (S200), we found a similar trend as S400, although the position difference was not as marked. The comparison of the mean difference between left and right hand position across the first and last five successful trials for all stiffness conditions (two-way mixed ANOVA) revealed a significant effect of Trial (F(1,21) = 28.05, *p* < 0.001). Post-hoc t-tests showed significant changes only in S400 and S200 conditions (*p* < 0.001 and *p* = 0.017, respectively), but not in S0 condition (**Fig. 3c**). We then quantified the temporal relation between the paired agents by computing the difference in time at which each agent moved past the 0.02-m distance from the starting position. A negative value would suggest temporal lead of the left agent. We again found that the left agent gradually increased the lead over the right agent only when the haptic feedback was enabled (S400 and S200). Two-way mixed ANOVA revealed a significant effect of Trial × Stiffness interaction (F(2,21) = 8.24, *p* = 0.0023). Post-hoc t-tests showed significant changes only in S400 and S200 conditions (*p* < 0.001 and *p* = 0.0025, respectively), but not in S0 condition (**Fig. 3d**). These results indicate that the stiffness of the dyadic interaction facilitated joint learning of a consistent spatiotemporal relation.

**Figure 3.**
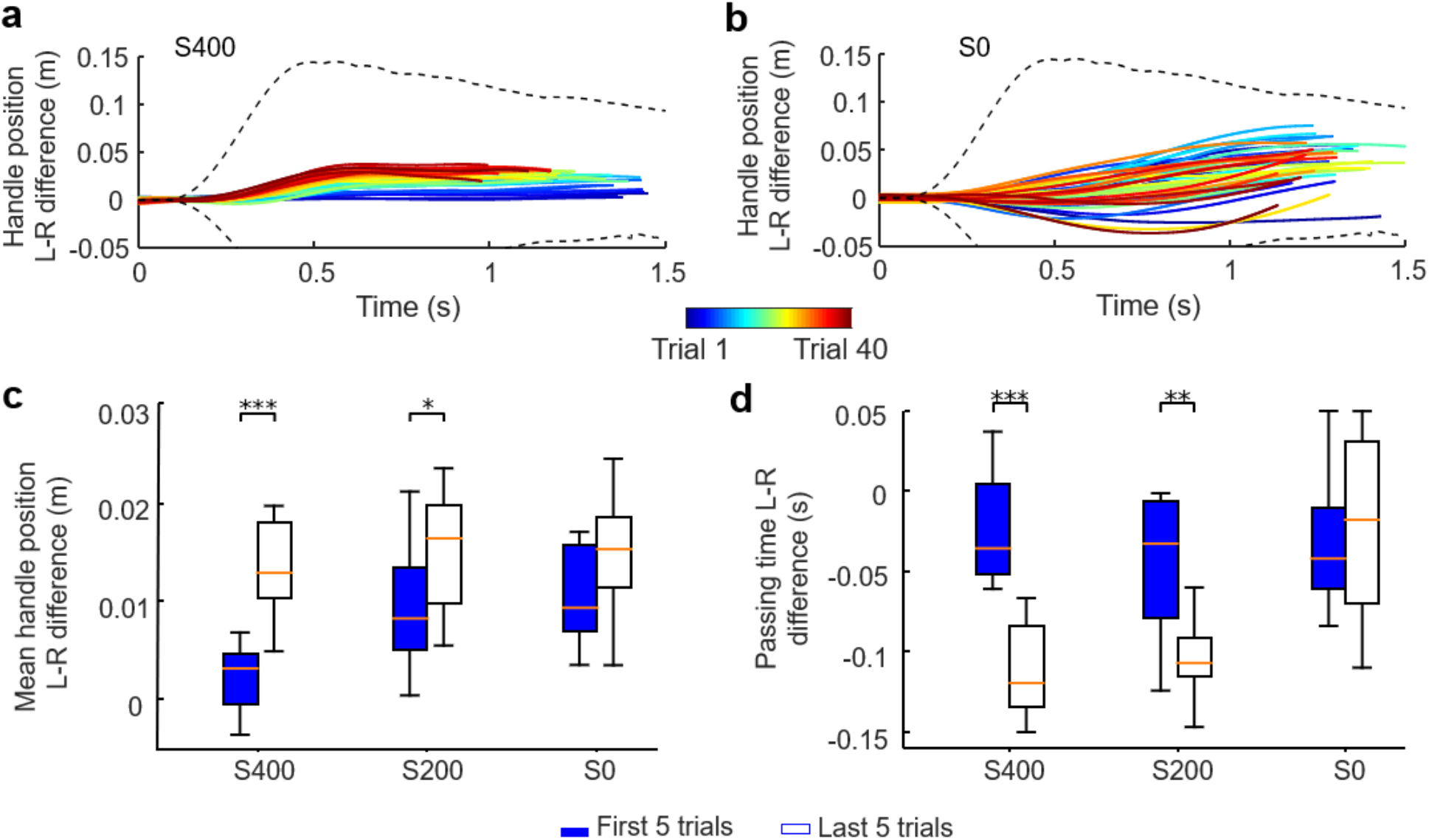
Changes of spatiotemporal coordination across successful trials. **a** Hand position difference from individual successful trials performed by a representative dyad in the S400 condition. Early trials are plotted as blue traces and subsequent trials are denoted by ‘warmer’ colors. Dashed lines represent boundaries of feasible strategies obtained from simulations. **b** Hand position difference from a dyad in the S0 condition are plotted in the same format as **a. c** Comparison of mean hand position difference between the first and last five successful trials. **d** Comparison of time at which left and right handles passed the 0.02-m distance from the starting point from the first and last five successful trials. (*, ** and *** denote p<0.05, 0.01 and 0.001, respectively). Rainbow-like color distributions can be seen in **a** but not in **b**, indicating the emergence of consistent co-adaptation only in the S400 condition and not in S0.

We then quantified the forces measured at the handles and how they changed across successful trials. We found that for both agents in the S400 condition, force magnitude gradually increased as their coordination strategy changed across trials, and the left agent used larger force than the right one (**Fig. 4a**). The contribution of two handles in S200 condition were characterized by a similar trend as the S400 condition, but no change in force was observed across trials. In contrast, there were no consistent difference in handle forces in the S0 condition between agents or trials. A three-way ANOVA confirmed this observations by revealing significant interactions (Trial x Stiffness: (F(2, 42) = 20.81, *p* < 0.001); Agent x Stiffness (F(2, 42) = 26.69, *p* < 0.001). Post-hoc t-tests revealed significant differences between first and last five trials for both agents only in S400 condition (*p* < 0.001), whereas both S400 and S200 conditions showed the left agent exerted larger force than the right agent (*p* < 0.001). Overall, these results suggest that dyads became less energy efficient in the higher stiffness conditions as participants gradually co-adapted towards new coordination strategies.

**Figure 4.**
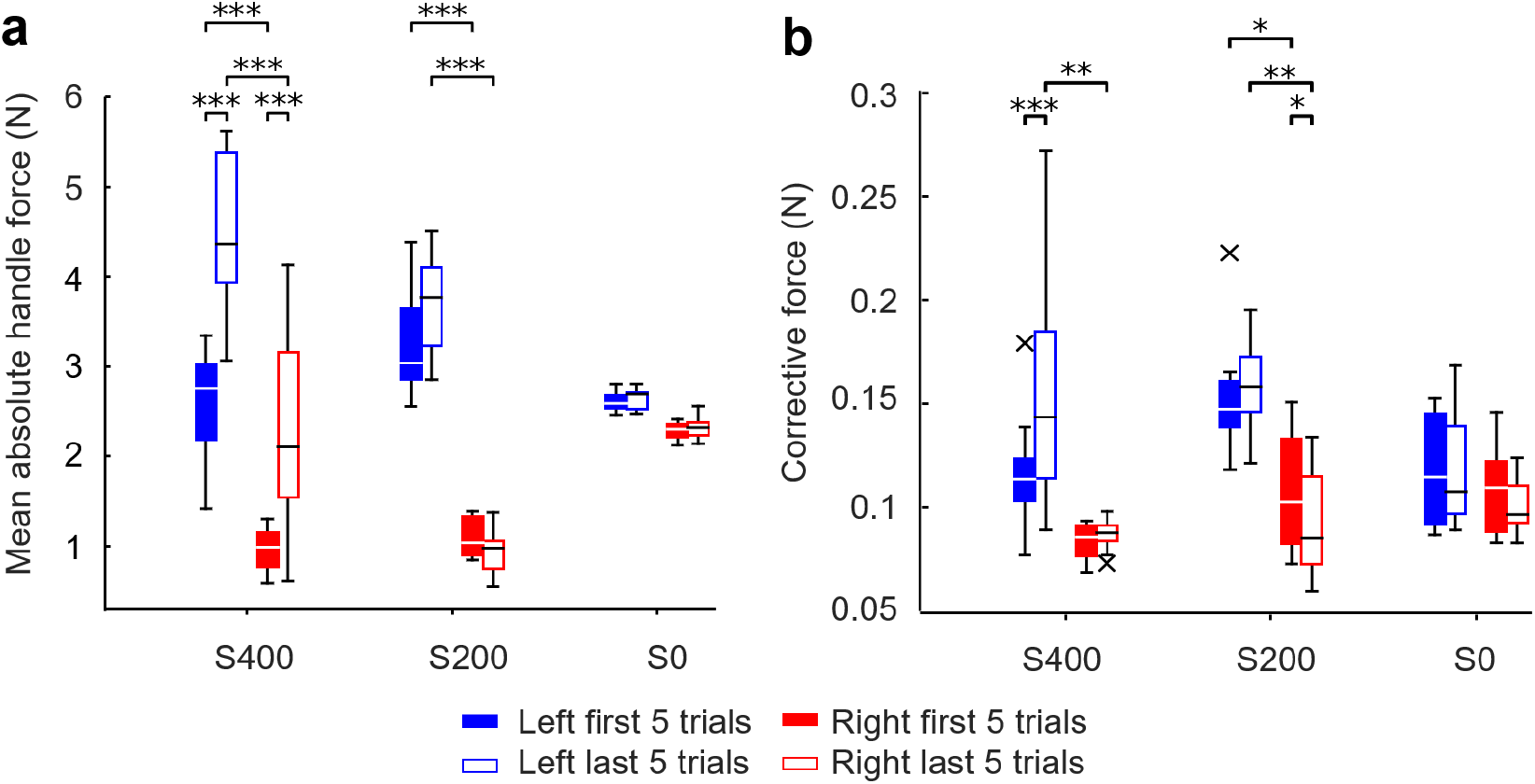
Changes of measured handle forces. **a** Comparison of mean absolute handle force between first and last five successful trials. **b** Comparison of corrective force between first and last five successful trials. Asterisks denote statistical significant differences between conditions (*, ** and *** denote p<0.05, 0.01 and 0.001, respectively). The ‘x’ symbols in the whisker plot denote outliers (1.5 interquartile range).

We also compared the corrective forces generated by each agent, measured as the deviation from the within-trial moving average (see **Methods**). We found that the leading (left) agent used more corrective forces. Moreover, the magnitude of the corrective forces only increased on later trials for the largest stiffness in the left agent, but not in the right one (**Fig. 4b**). A three-way ANOVA revealed multiple significant interactions (Trial x Stiffness: (F(2, 42) = 6.81, *p* = 0.003; Agent x Trial: (F(1, 42) = 11.13, *p* = 0.002. In the S400 condition, post-hoc t-tests revealed significant differences between the first and last five trials for left agents (*p* < 0.001), but not for right agents. Moreover, during the last five trials of S400 and S200 conditions, the left agents produced significantly larger corrective force than the right agents (*p* < 0.01 in both conditions). However, in the S0 condition no significant difference was found between agents or across trials. These results suggest that, as dyads co-adapted their coordination strategy across trials, the agents who led spatiotemporally were also more active in generating corrective actions.

What is the functional role of the systematic strategy exploration, i.e., converging to a consistent leader-follower relation, that emerged in the higher stiffness conditions? To answer this question, we quantified the stability of the beam as the maximum beam angle during the transport movement. Our task required the beam angle to be less than 1.15 degrees throughout the movement to be successful (see **Methods**). Therefore, a smaller maximum beam angle within a trial is considered to be indicative of greater stability. For the S400 condition, we observed a strong negative linear relation whereby the maximum beam angle decreased across successful trials (**Fig. 5a**). The S200 condition exhibited a similar, but weaker maximum beam angle reduction trend than S400. In contrast, the S0 condition did not exhibit consistent trial-to-trial changes in maximum beam angle (**Fig. 5b**). These observations were confirmed by a two-way repeated ANOVA revealing a significant Trial x Condition interaction when comparing first and last five trials across stiffness conditions (F(2, 21) = 17.43, p < 0.001; **Fig. 5c)**. Post-hoc tests revealed a significant difference only in the S400 condition (p < 0.001). These results suggest that the systematic adoption of new coordination strategies in the S400 condition might have enabled improvement of overall stability of the transport movement. To further illustrate the co-adaptation of the dyads, in **Figure 5d** we show how the coordination strategy (i.e., mean hand position difference) and the maximum beam angle evolved from the first five to the last five successull trials. It can be clearly observed that S400 dyads were able to converge towards optimal strategies, whereas S200 and S0 dyads did not.

**Figure 5.**
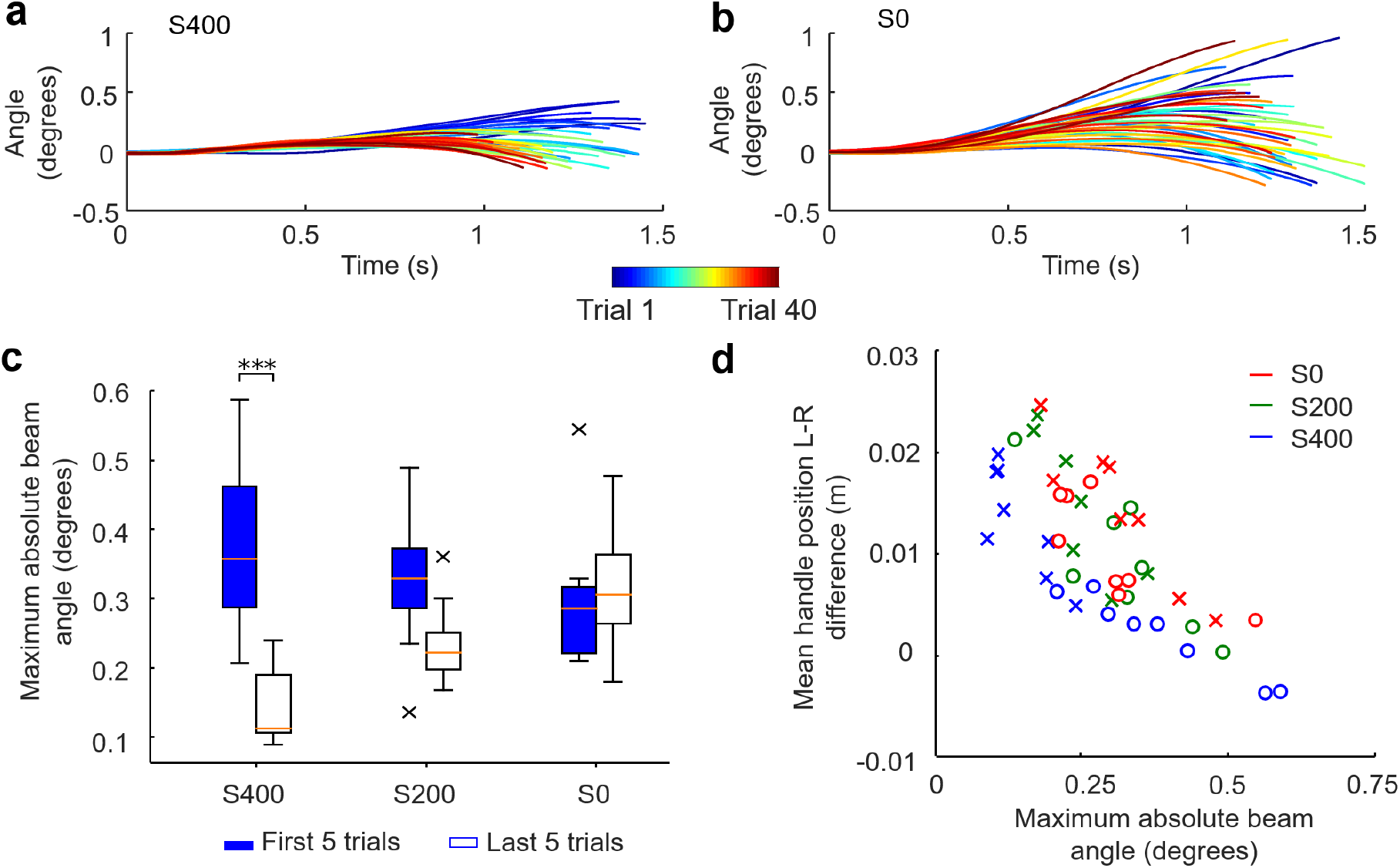
Changes of maximum beam angle across successful trials. **a** and **b** show maximum beam angle from individual successful trials performed by one dyad in the S400 and S0 conditions, respectively. Early trials are plotted as blue traces, and subsequent trials are denoted by ‘warmer’ colors. **c** Comparison between first and last five successful trials within each experimental condition (*** denote p < 0.001). The ‘x’ symbols in the whisker plot denote outliers (1.5 interquartile range). **d** Relationship between maximum absolute beam angle and mean hand position difference. Each data point denotes the average of the first and last five successful trials (circles and crosses, respectively) from all dyads. Blue, green, and red symbols denote data from S400, S200, and S0 conditions, respectively.

## DISCUSSION

In the present work we used a task (**Fig. 1**) that allowed a large family of successful solutions (**Supplementary Fig. 1**) which provide implicit gradients for optimizing dyadic motor behavior. We tested the hypothesis that the very existence of such optimization gradients would lead to gradual co-adaption, culminating with convergence to a consistent strategy. Importantly, we also hypothesized that such convergence would be stronger when veridical haptic information can be exchanged between the two agents. Our results support both hypotheses. Following a systematic exploration of the solution space, the experimental conditions characterized by haptic feedback (S400 and S200) led dyads to co-adapt and converge to similar solutions whereby one of the agents’ (left) movement spatio-temporally led the other agent’s (right) movement (**Fig. 3**). Moreover, the left agent exerted greater force and exhibited more corrective forces than the right agent (**Fig. 4**). These asymmetrical contributions are consistent with various notions of role specialization or leader-follower relation in human-human cooperation, e.g., spatial or temporal lead-lag, as well as differences in force contributions or corrective responses^9–16,25^. In contrast, when the haptic feedback was absent, dyads did not exhibit systematic co-adaptation. These findings, for the first time, reveal that consistent leader-follower relationships emerge from implicit haptic interactions.

To address the functional purpose of the leader-follower relation that spontaneously emerged, we analyzed three potential variables: movement time, the force exerted on each handle, and the maximum beam angle. Although movement time improved across trials for all stiffness conditions (**Supplementary Fig. 1**), it was not dependent on the availability of haptic feedback. Additionally, the exerted forces also increased across trials as the role-specialization emerged (**Fig. 4**), requiring agents (especially the left one) to increase energy expenditure. Therefore, it is unlikely that the main objective of co-adaptation we observed in conditions with haptic feedback was to achieve faster movements or miminze metablic cost. However, analysis of beam angle revealed that the largest stiffness condition enabled dyads to decrease the maximum rotation of the beam during movement (**Fig. 5**), which occurred despite the fact that dyads made no large error outside of the tolerance boundary. Maintaining a safety margin has been shown to be an important goal for individual human motor control, especially in grasping behaviors^26,27^, to account for uncertainty in the environment^28^. The present results suggest that both agents might have pursued the same objective of maximizing ‘safety margin’ for performing the task, thus enabling co-adaptation of motor coordination.

Importantly, our findings suggest that haptic feedback is necessary for co-adaptation to occur in our task. Why was visual feedback alone unable to mediate co-adaptation towards a consistent strategy? When an individual is required to adapt to a novel task dynamic, a systematic trial-to-trial change of strategy can only occur if they can infer the dynamics of the system they interact with^29–31^. Interestingly, it has been shown that vision alone could enable individuals to create internal representations of force fields^32,33^ or object properties^34,35^. However, in these studies the dynamics to be inferred was invariant across trials. In contrast, our task requires each agent to identify system dynamics that arises from both an object and another agent who also acts on the object. Therefore, in our task visual feedback may not provide sufficient information to enable a participant to infer the relative contribution of his or her own motor action to the beam’s movement, especially when considering that the hand position was hidden during the movement. It is therefore plausible that haptic feedback, especially in the high stiffness condition, provides important information about the consequence of one’s own actions, i.e., the magnitude and the direction of the beam movement with respect to one’s hand. Even though the importance of haptic feedback to infer the dynamics of a system that may consist of one or more human agents has been reported by previous work^23,36^, for the first time the present findings demonstrate its role for dyadic co-adaptation.

Although role specialization has been commonly demonstrated in previous work on dyadic motor coordination, consistent coordination patterns were not found across different dyads for a given task even when haptic feedback was presented^15,16,37^. It has been speculated that the dyad-specific coordination patterns may be determined by differences in the sensorimotor control capability (e.g., speed, accuracy, or strength) of the paired agents. For instance, agents with different reaction times and movement speed could lead to spatiotemporal asymmetry in joint reaching^21^. In contrast, we demonstrated that the same consistent leader-follower relation emerged across different dyads in our task. As noted in the **Introduction**, an important feature of our task is the sutble asymmetry in the mechanical moment arms that left and right agents used to move the beam. While it is important to point out that such asymmetry does not require a particular coordination pattern to perform the task successfully, our task inherently provides gradients for optimization as some strategies are associated with less energy expenditure whereas others enable better stability. This is different from previous work that used symmetrical dyadic tasks in which no clear directional advantage existed for agents’ asymmetrical contribution. Therefore, swapping the direction of asymmetrical role between agents in previous studies would not have significantly impacted task performance, i.e., either agent can perform any role, notwithstanding inter-agent differences in sensorimotor control capabilities. We conclude that the directional advantage introduced by context asymmetry is a critical factor for the emergence of leader-follower relations leading to a global solution for attaining a task goal. This conclusion is supported by findings from Takai and colleagues (personal communication) who found no dyadic coadaptation toward leader-follower relationships in a similar experimental setting where the task context was symmetrical. Our findings also suggest that co-adaptation of cooperating agents can exploit context asymmetry to jointly optimize a performance variable. Whether asymmetry in agents’ sensorimotor skill capabilities might be capable of generating the same phenomenon remains to be investigated.

The present results raise the question of whether the formation of leaders and followers in different contexts, i.e., groups of individuals or societies, might also be facilitated by asymmetrical contexts. Assuming the existence of a common goal bringing together multiple individuals, a subgroup of individuals (followers) might become attracted towards an individual (leader) because of asymmetrical contexts, including knowledge, educational background, and access to resources, e.g., wealth, marketing, etc. An additional analogy between our task and large-scale cooperation is the critical role of effective communication between group members^38^. In our task, removal of haptic feedback interfered with the emergence of leader-follower relations and their convergence to a stable strategy. Thus, communication is necessary to enable cooperating agents to infer their own contributions to a given ‘big picture’, shared goal.

This study is the first to report the gradual emergence of, and convergence to, consistent asymmetrical roles in dyads. This role specialization appears to have emerged through similar mechanisms proposed for the regulation of human cooperation, i.e., collective intentionality^1,3^, which includes a common understanding of interaction rules followed by both agents in a dyad with respect to the asymmetrical task context through effective communication channels. Our findings provide solid neuroscientific support to the hypothesis that “leaders are made by their assigned initial roles both explicitly or implicitly”, or acquisition rather than innate nature for the generation mechanism. Although role sharing appears to emerge to optimize motor performance and, possibly, cognitive loads of the cooperating agents^2^, future work is needed to identify the neural mechanisms underlying dyadic co-adaptation.

## Acknowledgments

This research was supported by the Commissioned Research of NICT, AMED under Grant Number JP21he2202005, Innovative Science and Technology Initiative for Security Grant Number JPJ004596, ATLA, Japan, JST [Moonshot R&D][Grant Number JPMJMS2034], and JSPS KAKENHI Grant Number JP20K20263.

## METHODS

### Participants

48 healthy right-handed subjects participated in this study. All participants provided written informed consent before participation. The study protocol was approved by the ATR Review Board Ethics Committee.

### Experimental apparatus

In dyadic conditions (see below), each subject used a robotic manipulandum, a twin visuomotor and haptic interface system^39^, to control a simulated virtual beam (**Fig. 1a**). Each robot’s arms consist of parallel links which are driven by electric motors placed under the display board on which the task was visually rendered by a projector. The handles of the manipulanda are aero-magnetically floated on the support table to minimize friction, such that they move freely on the flat surface of the support table. In the present study, the handle was programmed to move only in forward-backward dimension (y; **Fig. 1b**). The forces exerted by a participant on the robot handle were measured by a six-axis force/torque sensor. Data were collected at a sampling rate of 2 kHz.

Participants sat on a reclining adjustable chair and wore a seat belt. We set the chair’s height so as to align the shoulder-arm-hand with the handle’s height on the horizontal plane. We positioned each participant by having his/her shoulder 45 cm away from the origin of the hand position. Each participant’s forearm rested on a cuff that was mechanically supported on the same horizontal plane. Therefore, participants were not required to hold their arm against gravity. For safety reasons, movement of the Tvins could be stopped if one of the following criteria was met: when the emergency hand switch was pressed by the participant, when the force/torque sensor measured force greater than 20 N, or when two participants’ hands were 20 mm away from each other on the y-axis. Each participant grasped the robot handle under the display board with his/her right hand. Additionally, we placed a partition to prevent participants from seeing each other. Participants wore earplugs and soundproof earmuffs to mute the robot sound.

### Virtual beam model

In all experiments, subjects held the robotic handles to move a virtual beam in the horizontal plane with a displayed length of 10 cm (**Fig. 1b**). The motion is constrained in y-direction by a virtual frictionless rail, so that participants cannot move in x-direction. The underlying dynamics of the beam motion were simulated by using the following translation and rotation dynamics (Eq. 1 and 2, respectively):

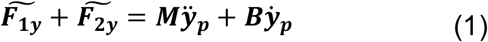

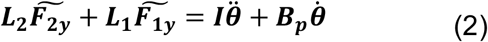

Here the motion of the beam was driven by two virtual input forces 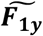 and 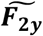 generated by the motion of the left and right hands with respect to the virtual beam (Eq. 3):

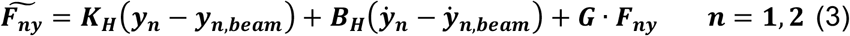

In these equations, ***y***_***p***_ and ***θ*** are the pivot position and rotation of the beam, ***y***_***n***_ and ***y***_***n***,***beam***_ are position of a hand and its corresponding virtual attach point on the beam, respectively. In our experiments, the distances (i.e., moment arms) from the left and right virtual attach points to the pivot location are ***L***_**1**_ and ***L***_**1**_, respectively. This design resulted in an asymmetric configuration in which the left virtual attach point has a smaller moment arm than the right one (**Fig. 1c**). In contrast to the dynamic model, the visual rendering of the beam only displays the portion of the beam between the two virtual attach points, which was scaled down as 10% in the x-direction to fit to the visual display.

To generate haptic feedback forces ***F***_***ny***,***fb***_ delivered to the corresponding hands, we programmed a haptic feedback model that computes the interaction force between the beam and participant’s hand with a spring-damper mechanism (**Fig. 1c**). X-directional feedback by the beam movement is not simulated in this study. By changing the stiffness coefficients of the spring, we could vary the transmission of force feedback from the beam to each participant as follows:

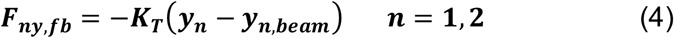

Parameters of equations (1)-(4) are shown in Table 1. It is important to point out that ***K***_***T***_ represents the stifness of the inter-subject physical coupling, which is the primary experimental factor we systematically investigated in this study. Specifically, we tested three stifness conditions (**K**_***T***_: 400, 200, and 0 N/m). We refer to these conditions as S400, S200 and S0, respectively.

**Table 1.** Coefficients of the virtual beam

| Coefficients | Value |
| --- | --- |
| $M$ Weight of the virtual beam [kg] | 1 |
| $I$ Moment of inertia of the virtual beam [kg m<sup>2</sup>] | 33.3 |
| $L_1$ Distance between left virtual beam attachment point and pivot [m] | 9 |
| $L_2$ Distance between right virtual beam attachment point and pivot [m] | 10 |
| $B$ Virtual beam translational coefficient [Ns/m] | 0.1 |
| $B_p$ Virtual beam rotational coefficient [kg/s] | 0.1 |
| $K_H$ Force input spring stiffness [N/m] | 100 |
| $B_H$ Force input damper coefficient [Ns/m] | 5 |
| $G$ Force input gain [-] | 0.1 |
| $K_T$ Force feedback spring stiffness [N/m] | Condition dependent |

### Experimental Protocol

Subjects were not informed that the task was to be performed in a collaborative fashion in the dyadic conditions, as they only interacted with one side of the virtual beam. The virtual beam, the start and target areas, and the tilt visual indicator were displayed on the table (**Fig. 1a**,**b**). The table was opaque, and therefore participants could not see their hand. The virtual beam was displayed as a thin white line, and the tilt visual indicator was a yellow circle sliding along the beam (**Fig. 1b**). The tilt indicator (yellow circle) slides from the center to the the higher of the two beam’s ends when it tilts from the horizontal to the maximum allowable angle. The circle behaves like an air bubble in a level scale. Therefore, when tilt occurs, participant could see both the angle of the beam and the displacement of the tilt indicator. The sensitivity of the tilt indicator was set to 0.4 cm/degree in the visualization (corresponding to approximately 43.6 cm/degree in the simulation model). Start and target areas of both ends of the beam had visual radii of 0.5 and 1 cm, respectively. Both participants’ target areas were displayed 10 cm away from the start area. Subjects were instructed that the handle moves only in y-direction towards the goal area.

Subjects were instructed to start the movement as soon as they saw the start area color change (movement start cue). After the beam reached the target, subjects were asked to relax and let the robotic manipulandum slowly move their hands back to the vicinity of the start area. Subjects had to re-enter the start area by themselves. A few seconds after both subjects re-entered the start area, a new trial was started.

Computation of movement time started when the hand of one of the participants left the start area. Dyads had to reach the target position with a minimum movement duration of 1 second and not longer than 2 seconds from movement start. When either criterion was not met, the trial was aborted and repeated. During the movement, participants were instructed that the beam angle should not exceed ±1.15 degrees. A trial was defined as successful when dyads moved the beam to the target by complying with both of the above spatial and temporal requirements. Movement time and maximum tilt angle were computed at the end of each trial and fed back to the participant on the visual display. If both spatial and temporal requirements were met, the message “Good” was displayed. Otherwise, if the movement time was shorter than 1 second or longer than 2 seconds, the message “short” or “long” was displayed, respectively. Additionally, “NG” was displayed when the tilt angle exceeded the maximum allowable value. Data collection was terminated when subjects performed 40 successful trials.

### Simulation of Coordination Strategies

The coordination strategies afforded by the experimental task were simulated by assuming a minimum jerk model for hand kinematics:

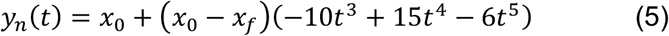

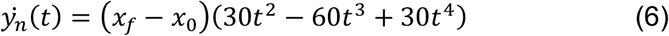

where the initial and final velocity, 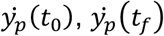 acceleration, 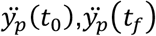, and the initial position, *x*_0_, are zero *x*_*f*_ is the final position. This kinematic model was scaled with a range of movement times (0.8 – 2.2 s) and final positions (0.15 - 0.21 m) to generate a wide range of kinematic profiles which were used to drive the beam simulation (Eq. 1-3). The output of these simulations was evaluated the same way as the experimental results by testing against the spatial and temporal requirements, and the successful simulations were considered as a feasible coordination strategy. The feasible strategy space was considered to range from the minimum and maximum mean hand position differences of these successful simulations (i.e., [-0.01 m, 0.06 m]). Note that the simulation did not involve any haptic feedback term, therefore the movement kinematics of the two hands were independent (as in S0 condition). Any feasible solution defined by this method would be a feasible solution in terms of hand kinematics for all stifness conditions.

### Corrective forces

The corrective force *F*_*nc,i*_ of each handle (*n* = 1, 2) at each trial (*i*) is derived by taking the average of the absolute difference between the measured handle force and the smoothed measured handle force (simple moving average, SMA) using the following equation,

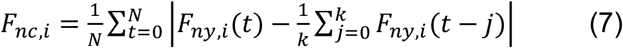

Where *F*_*ny,i*_(t) is the measured handle force at time *t*, and *N* is the number of samples within the experimental time (*t*_0_, to *t*_*f*_). We set the sliding window length *k* as 0.2 seconds and used the MATLAB function *movmean*() for the SMA calculation.

