## Supplementary material for "Leaders are made: Learning acquisition of consistent leader-follower relationships depends on implicit haptic interactions"

***1. Simulation of feasible space of dyadic motor interactions.*** Computer simulations using minimum jerk hand kinematics (**Supplementary Fig. 1a**) suggested that the range of mean hand position difference (Left – Right) to comply with the task's spatial and temporal requirements is -0.02m to 0.06 m. The beam angle during these coordinated movement is shown in **Supplementary Fig. 1b**. Note that some coordination strategies can lead to a smaller peak beam angle, thus denoting a potential optimization gradient.

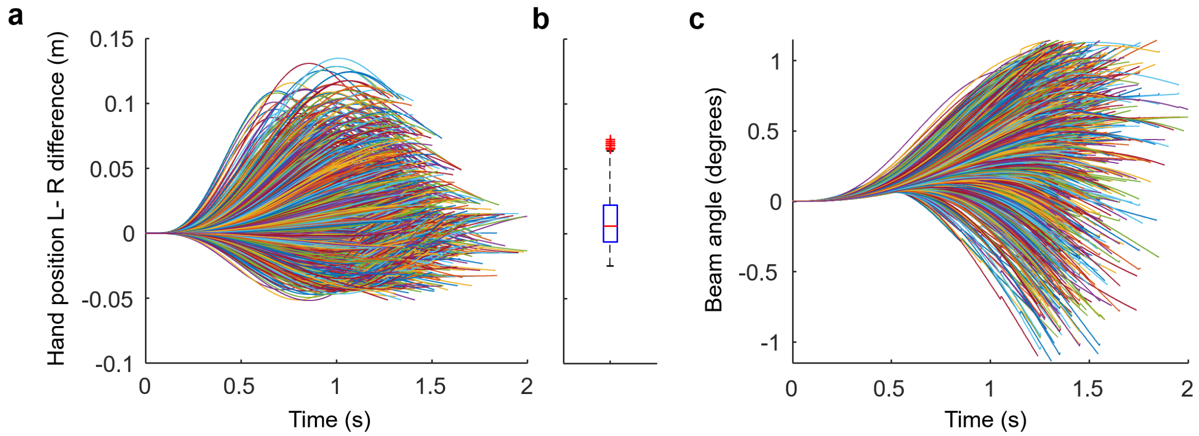


**Supplementary Figure 1. Simulation of feasible coordination strategies.** **a** Hand position differences (Left – Right) used in successful trial simulations and the boundaries of hand position strategies. **b** Distribution of mean hand position differences (Left – Right) of feasible coordination strategies. **c** Beam angle during successful trials. Each colored line in **a** and **c** denote one successful trial simulation.

***2. Total Trial and Movement speed.*** Participants were able to accomplish 40 successful trials across all stiffness conditions with only a few failed trials (significant effect of Condition, Kruskal-Wallis test, $\chi^2 = 8.2$, p = 0.016; **Supplementary Fig. 2a**). Movement speed significantly increased across successful trials in all conditions (two-way mixed ANOVA, significant main effect of Trial on movement speed: $F(1,21) = 21.54$, $p < 0.001$, **Supplementary Fig. 2b**).

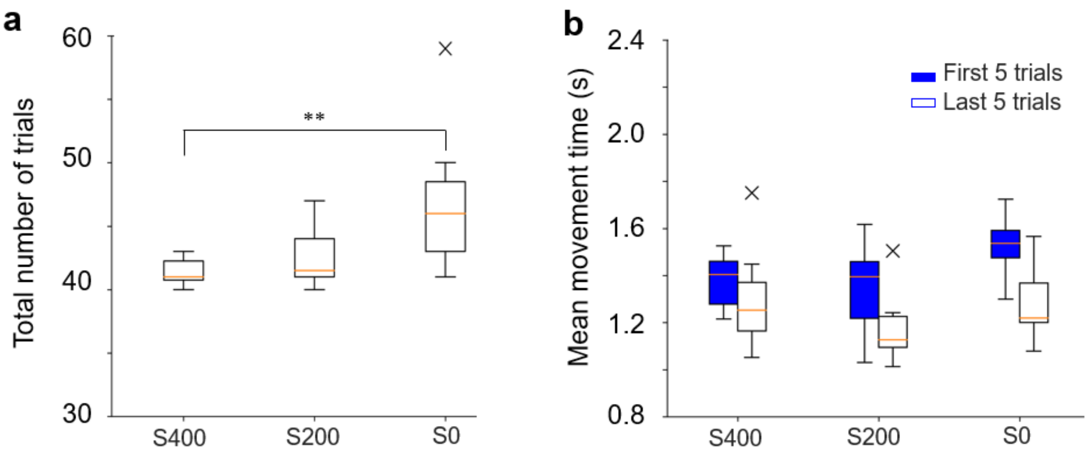

**Supplementary Figure 2. a** Total number of trials required to accomplish 40 successful trials across stiffness conditions. Asterisks denote statistically significant differences between conditions (** denote p<0.01). **b** Comparison of movement time between the first and last five successful trials within each stiffness condition. The '×' symbols in the whisker plot denote outliers (1.5 interquartile range).

***3. Dyadic strategy optimization***. Simulation of the effects of feasible dyadic interactions on maximum beam angle showed that the optimization landscape is approximately '<' shaped with a global minimum (**Supplementary Fig. 4a**). The experimental results show that only the S400 dyads could learn to systematically operate in this space (blue crosses, **Supplementary Fig. 4b**).

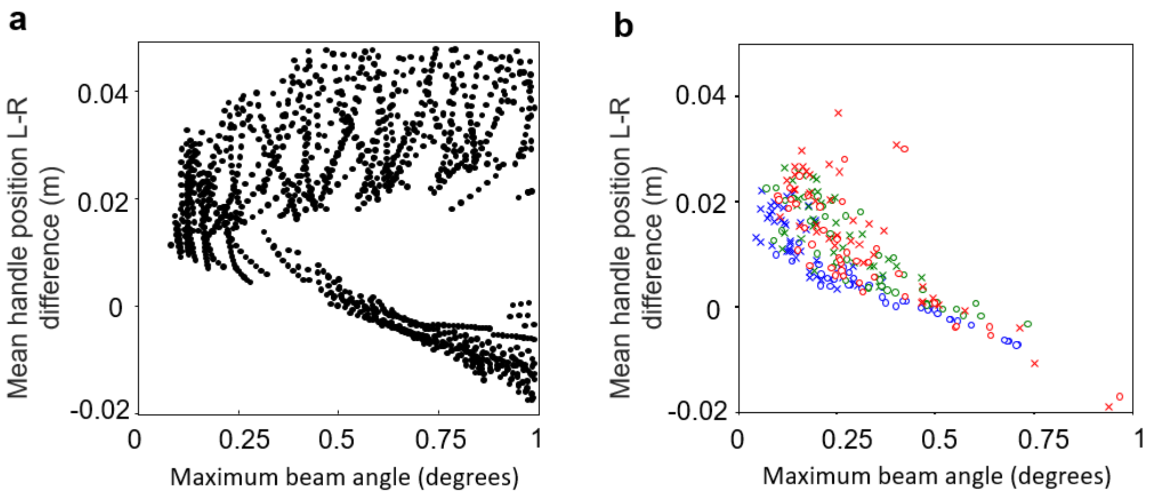


**Supplementary Figure 3. Motor coordination optimization. a** Simulated results (sampled from 1500 successful simulations) of successful relations between mean hand position difference and maximum beam angle. **b** Results from first and last five successful trials (dots and crosses, respectively) from all dyads. Blue, green, and red symbols denote data from S400, S200 and S0 conditions, respectively.
